# An arachnid’s guide to be an ant: morphological and behavioural mimicry in ant-mimicking spiders

**DOI:** 10.1101/2021.10.30.466574

**Authors:** Nimish Subramaniam, Krishnapriya Tamma, Divya Uma

**Affiliations:** School Arts and Sciences, Azim Premji University, Bangalore

**Keywords:** Myrmaplata, ant mimicry, morphometrics, shape analysis, myrmecomorphy, movement

## Abstract

Batesian mimicry imposes several challenges to mimics and evokes adaptations in multiple sensory modalities. Myrmecomorphy, morphological and behavioral resemblance to ants, is seen in over 2000 arthropod species. Ant-like resemblance is observed in at least 13 spider families despite spiders having a distinct body plan compared to ants. Quantifying the extent to which spiders’ shape, size, and behavior resemble model ants will allow us to comprehend the evolutionary pressures that have facilitated myrmecomorphy. *Myrmaplata plataleoides* are ‘accurate’ mimics of the weaver ants, *Oecophylla smaragdina*. In this study, we quantify the speed of movement of model, mimic, and non-mimetic jumping spiders. We use traditional and geometric morphometrics to quantify traits such as foreleg and hindleg size, body shape between the model ant, mimic, and non-mimics. Our results suggest that while the mimics closely resemble the model ants in speed of movement, they occupy an intermediate morphological space compared to the model ants and non-mimics. We suggest that ant-mimicking spiders are better at mimicking ants’ locomotory movement than morphology and overall body shape. Our study provides a framework to understand the multimodal nature of mimicry and helps discern the relative contributions of such traits that drive mimetic accuracy in ant-mimicking spiders.

## Introduction

Batesian mimicry, where a palatable, undefended species mimics an unpalatable, noxious model to avoid predation is a fascinating example of adaptive evolution (Bates 1862, Pfennig et al. 2001, Ruxton et al. 2004). Such mimicry poses several challenges to the mimics, and often elicits multimodal adaptations (Ruxton et al. 2004, Uma et al. 2013, Rowe & Halpin 2013, Pekár et al. 2017). Morphological and behavioral resemblance to ants (myrmecomorphy) is a well-known example of Batesian mimicry (Cushing 1997, Nelson et al. 2005, Nelson & Jackson 2006, Huang et al. 2011, Durkee et al. 2011). Because of their aggressive tendencies and communal defence that deter many generalist predators, ants serve as model organisms for a number of arthropods (McIver & Stonedahl 1993). In fact, over 2000 species across 54 families of insects and spiders have evolved morphological and behavioural resemblance towards ants (McIver & Stonedhal 1993). Since ants do not have strong aposematic colouration, ant-mimics resemble ants in locomotory behavior and the general body plan to avoid visually-oriented predators (Pekar & Kral 2002, Nelson & Card 2016). For example, ant-mimicking spiders derive protection from predators such as jumping spiders, mantids and mud-dauber wasps (Nelson & Jackson 2006, Durkee et al. 2011, Huang et al. 2011, Uma et al. 2013, Nelson & Card 2016, Ramesh et al. 2016).

Ants (subphyla: Hymenoptera) and spiders (subphyla: Chelicerata) belong to different classes that diverged approximately 540-600 mya (Regier et al. 2005). Ants and spiders have very different body plans: Ants have a slender shape, a distinct head, thorax, and abdomen, 6 legs, and a pair of antennae. On the other hand, spiders have a much more oval shape, fused head and thorax (cephalothorax), 8 legs, two extra leg appendages (metatarsus and patella) compared to ants, and no antenna. Mechanistically, ants use flexor-extensor muscles to move their legs, but spiders lack extensor muscles in major leg joints. They instead use hydraulic pressure generated in their bodies to move their legs (Kropf 2013, Hill 2018). In spite of these differences, ant-like resemblance can be seen in 13 spider families (Cushing 1997). How do spiders that mimic ants overcome these differences, given that their body plan is evolutionarily distinct from that of ants?

Ant-mimicking spiders have a constricted cephalothorax, elongated pedicel, a slender abdomen, and coloration with hairs and scales which helps them achieve an ant-like body form. Behavioral modifications include walking in a zig-zag motion, bobbing their abdomen, and ‘antennal-illusion’ by waving of the first pair of legs in air (McIver & Stonedhal 1993, Cushing 1997, Ceccarelli 2008, Shamble et al. 2017).

The extent to which ant-mimicking spiders resemble ants is constrained by predation pressure and several other factors. Slender body shape and thin legs may constrain accurate mimics in their jumping and prey capturing abilities (Hashimoto et al. 2020), while constricted abdomen may reduce fecundity (McIver & Stonedhal 1993). Developmental, genetic and life history parameters also impose constraints (Pekár & Jarab 2011, McClean et al. 2019). Quantifying the extent to which spiders’ body form and behavior is modified to resemble ants will help elucidate the evolutionary pressures that have facilitated myrmecomorphy.

Recent studies have measured various traits such as body shape, size, color, movement, and walking trajectories that contribute to mimicry. (Shamble et al. 2017, Hashimoto et al. 2020, Pekár et al. 2020, McLean & Herberstein 2021). However, these studies have not compared all three players (model, mimic, and non-mimic) in the mimetic system, nor have they included multiple traits that contribute to mimicry (but see Kelly et al. 2021). For example, Hashimoto et al. (2020) compare various non-mimetic jumping spider species to ant-mimicking jumping spiders in the genus *Myrmarachne*, and show that they occupy distinct morphometric space. However, they did not include ants in this comparison. McLean and Herberstein (2021) include mimics, non-mimics and ants and show that mimetic shape is intermediate to that of ants and non-mimics, but they did not include size measurements. Size considerations, particularly that of the legs, is important, as they influence jumping distance, speed, and trajectories in mimics and non-mimics (Hill 2018, Shamble et al. 2017). Expanding on previous studies, here we include both size and shape measurements along with locomotory behaviour of mimics, non-mimics and ants to understand the relative extent to which they contribute to mimetic accuracy.

*Myrmaplata plataleoides* (Salticidae) are considered ‘accurate’ morphological mimics of the weaver ants, *Oecophylla smaragdina* (Formicidae) (Edmunds 2006). This mimic and model system (Fig 1), found across Australia and Asia, has been extensively studied. *M. plataleoides* gains protection from visually-oriented predators such as mantids and jumping spiders (Ramesh et al. 2016, Vijayan et al. 2021). In this study, we investigate the extent to which mimics resemble their model ants compared to non-mimetic jumping spiders. Specifically, we compare a) behavioural traits such as the speed of movement, and b) morphological traits such as leg size, body shape, between model, mimic and non-mimic spiders.

**Fig.1.**
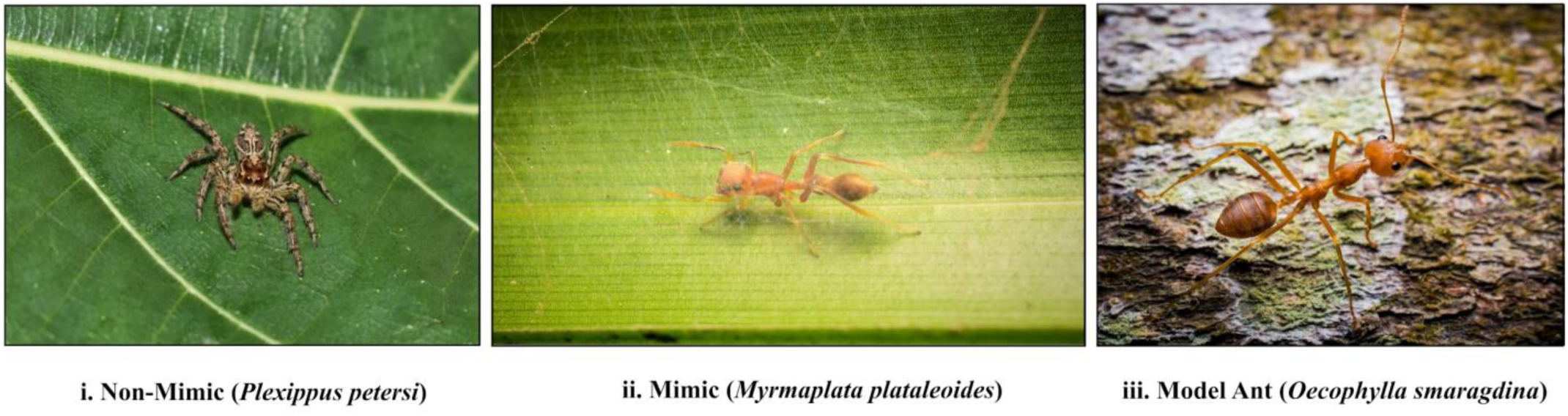
Non-mimic, ant-mimic and model ant used in the study. Representative species of non-mimetic jumping spider is given here. Sizes of the animals are not to scale. Photo credits: non-mimic: Nimish Subramaniam; mimic and model ant:Samuel John

**Fig.2.**
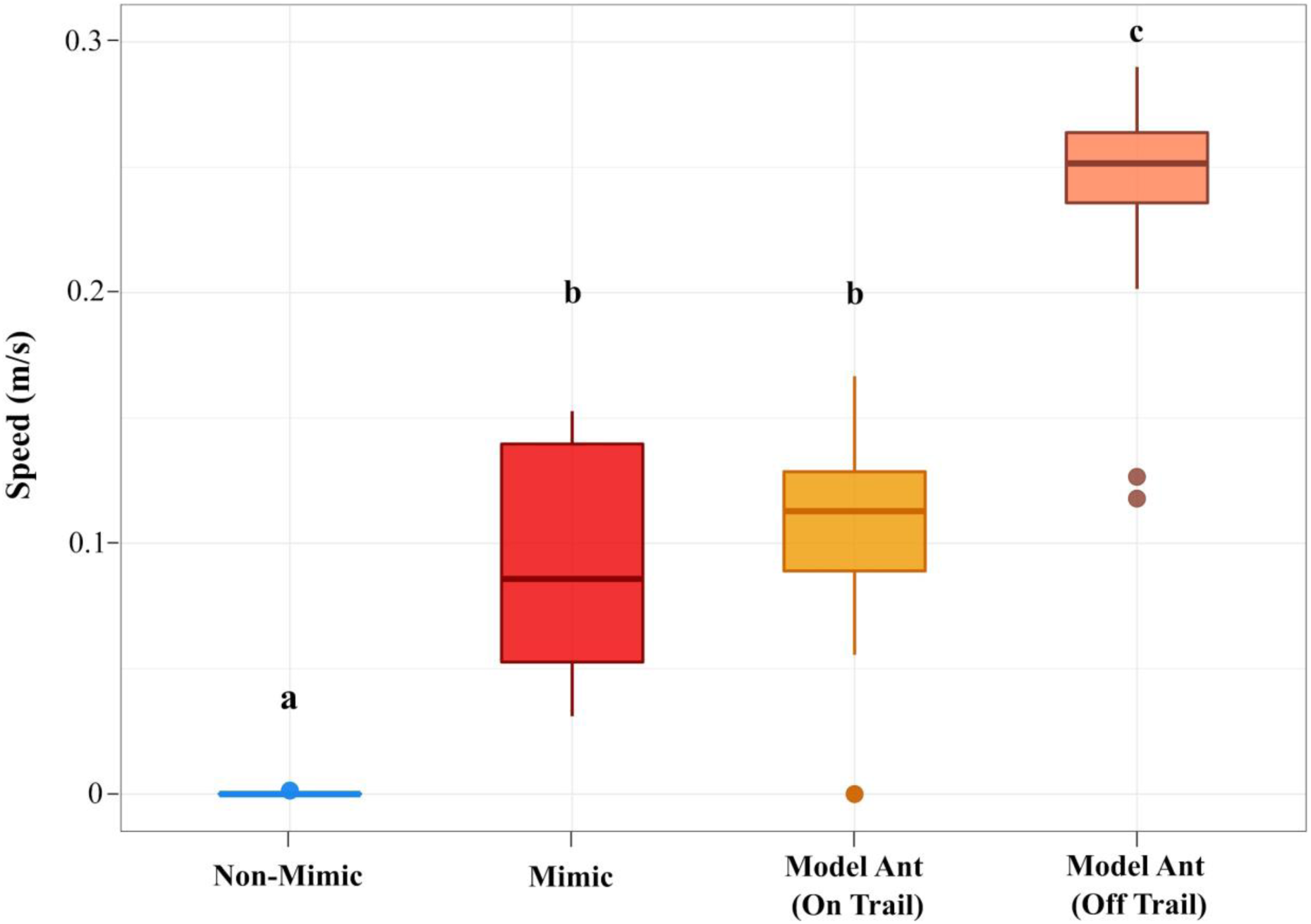
Speed of movement of non-mimics, mimics and model ants. All groups were significantly different from each other (Kruskal Wallis χ2 = 40.132, df = 3, p<0.001). Median speed of mimics is similar to that of a model ants following a pheromone trail (Bonferroni-Dunn test, p=1) and is significantly different from that of non-mimics (Bonferroni-Dunn test, p<0.005) and model ants walking in the absence of a pheromone trail (Bonferroni-Dunn test, p<0.005). Letters a, b and c denote significantly different values.

## Methods

### a) Study animals

Adult and subadult female *Myrmaplata plataleoides* (henceforth mimics), and various non-mimetic jumping spider species (henceforth non-mimics) were collected in and around Bengaluru, India between January 2018 and July 2019. Non-mimics were collected from the same habitat as that of mimics. They were housed in Azim Premji University in individual plastic containers with leaves and twigs and a moist cotton ball, and were maintained under a 12 hour day-night cycle. They were fed fruit flies (*Drosophila sp*.) every alternate day. Major workers of weaver ants, *Oecophylla smaragdina* (henceforth model ants) were collected on the day of the experiment near Sarjapura village, Bengaluru, India.

### b) Speed of movement

We compared the speed of movement between mimics, model ants and one species of non-mimetic spider (*Plexippus petersi*). Videos of mimics (N=16) and non-mimics (N=13) were taken as they moved freely on a featureless 40 cm X 20 cm arena. A Sony HDR-PJ410 handycam, placed perpendicular to the arena, was used to film their movements at 25fps. For each trial, an individual (mimic or non-mimic) was released at one end of the arena, and recorded as it moved to the other end. Trials were deemed successful only if individuals reached the other end without leaving the video frame in between. Individuals that failed to do so were subjected to a retrial after a period of one hour. The arena was wiped with ethanol between every subsequent trial to remove any chemical cues or silk deposited by the study animals. Animals were released back to the site of collection once trials were concluded.

As weaver ants are known to follow a pheromone trail, the speed of model ants was analyzed on and off a trail. Videos of individual ants (N=14) that did not follow a trail were recorded in the same manner as that of mimics and non-mimics described earlier. For the pheromone trail following ants (N=12), the movement of worker individuals going from the base of a mango tree trunk towards their nest was recorded.

We analysed speeds (by using TrajR) (McLean & Skowron Volponi 2018) of mimics, non-mimics and model ants based on their trajectories obtained from Tracker (Brown 2008). Kruskal-Wallis test was used for multiple comparisons, and Dunn’s test with Bonferroni corrections was used for post-hoc analysis.

### c) Leg size

We used model ants from two different colonies (colony 1, N=9; colony 2, N=18), non-mimics belonging to different salticid genera (*Plexippus sp*., N = 11; *Hasarius sp*., N=1; *Hyllus sp*., N=1, unknown, N=6), and mimics (N=10) to quantify leg morphometrics.

We first compared all eight legs of mimics and non-mimics based on the length: width measurements of their femur, patella, tibia, metatarsus and tarsus (Fig. 3A).

**Fig.3.**
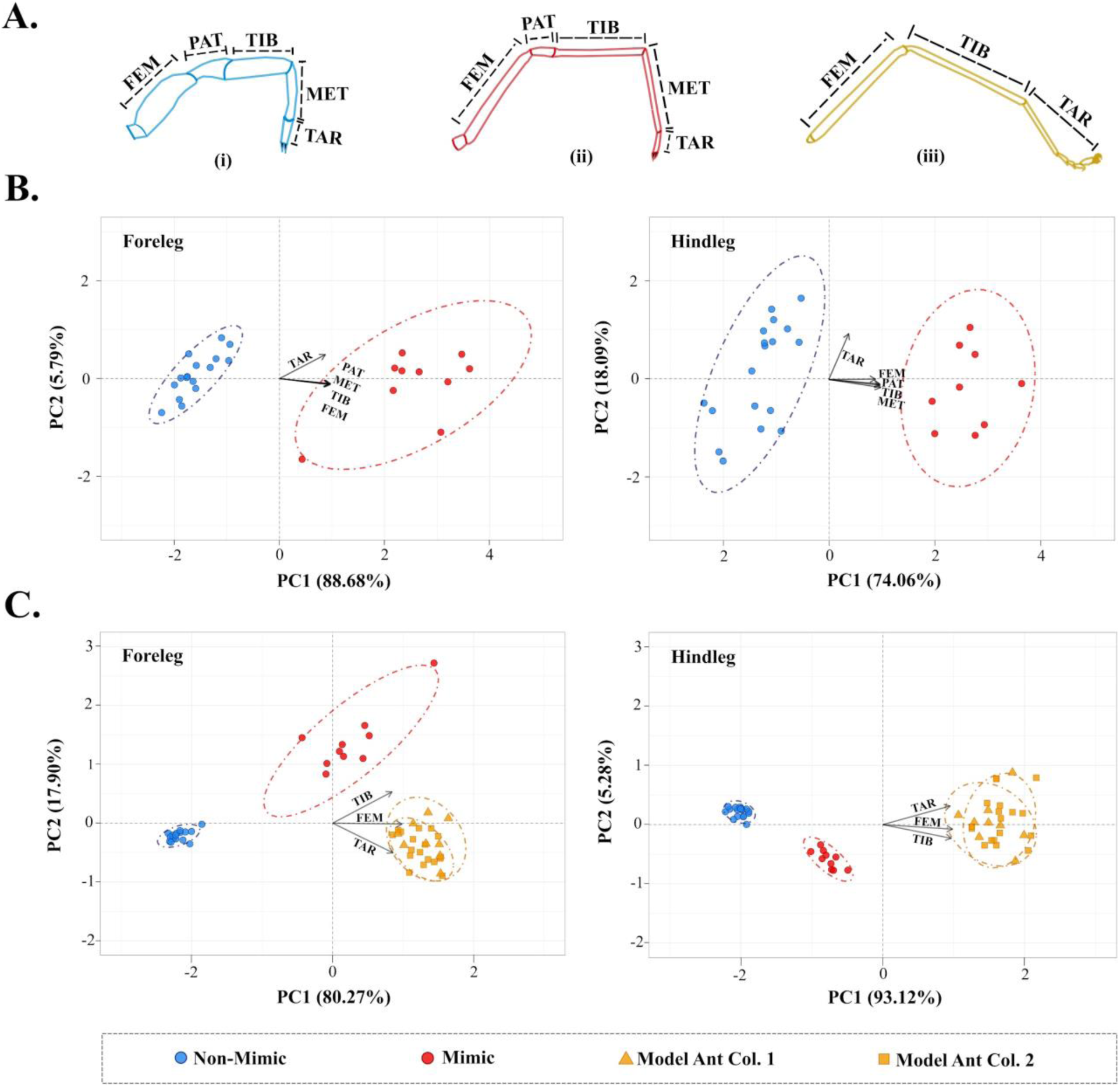
PCA of leg size of non-mimics, mimics and model ants. (A) Illustration of leg appendages of (i) non-mimics, (ii) mimics and (iii) model ants. (B) PCA results quantifying differences in the size of right foreleg and hindleg of mimics and non-mimics based on the length: width ratio of femur (FEM), patella (PAT), tibia (TIB), metatarsus (MET) and tarsus (TAR). Non-mimics and mimics show clear separation along the PC1 axis which accounts for 88.68% and 74.06% of the observed variation in foreleg and hindleg size respectively. (C) PCA results based on length: width ratio of leg appendages common to non-mimics, mimics and model ants viz. femur (FEM), tibia (TIB) and tarsus (TAR). Mimics occupy an intermediate position between non-mimics and model ants along the PC1 axis. For B and C, each data point represents measurements taken from the leg of an individual.

For comparing mimics, non-mimics and model ants, only leg I (henceforth ‘foreleg’) and leg IV (henceforth ‘hindleg’) were chosen as they are functionally important for ‘antennal illusion’ and jumping respectively. As ants lack patella and metatarsus, we performed a separate analysis in which we only included measurements of femur, tibia and tarsus, which are leg appendages common to mimics, non-mimics and model ants (Fig. 3A). Coxa and trochanter were excluded from both the analyses due to their small size, which makes them difficult to measure accurately.

Individual legs of specimens of comparable size (average body length ± SD: non-mimics: 7.19 ± 1.04 mm; mimics: 6.86 ± 0.73 mm; model ants colony 1: 8.13 ± 0.86 mm; model ants colony 2 - 9.57 ± 0.54 mm) were dissected at the coxa-trochanter joint and were photographed using a Laben STZ-450T microscope (with Tcapture) with a reference scale. Some specimens were examined during COVID-19 pandemic induced lockdown and owing to inaccessibility of lab resources, a Redmi Note 9 smartphone coupled with an iVoltaa 20X clip-on macro lens was used for imaging. The use of a mobile camera did not affect the quality of the images and required measurements could be taken accurately. Images were then analyzed using *ImageJ* (Schneider et al. 2012). We performed principal component analysis on the ratio of length: width values for aforementioned appendages of each individual leg using the R package *FactoMineR* (Le et al. 2008).

### d) Shape analysis

For landmark based analysis, images of the ventral side of mimics (N=11) and non-mimics (*Plexippus sp*. N=11) were taken under the Laben STZ-450T microscope. Nine landmarks were placed on each specimen, eight corresponding to the point where individual coxae fuse with the cephalothorax, and one placed at the base of the pedicel (Fig. 4A). Landmarks were digitized using *tpsDig* ver. 2.31 (Rohlf 2015, 2017) and the raw co-ordinates were superimposed using a generalized procrustes analysis (GPA) which removes the effect of size (Rohlf & Slice 1990, Lawing & Polly 2010) (Fig. 4B) using the R package *geomorph* (Adams & Otárola-Castillo 2013). We used relative warp analysis to calculate differences in the shape of these superimposed configurations. We did not include model ants as landmark based analyses can only be performed on homologous features of organisms.

**Fig. 4.**
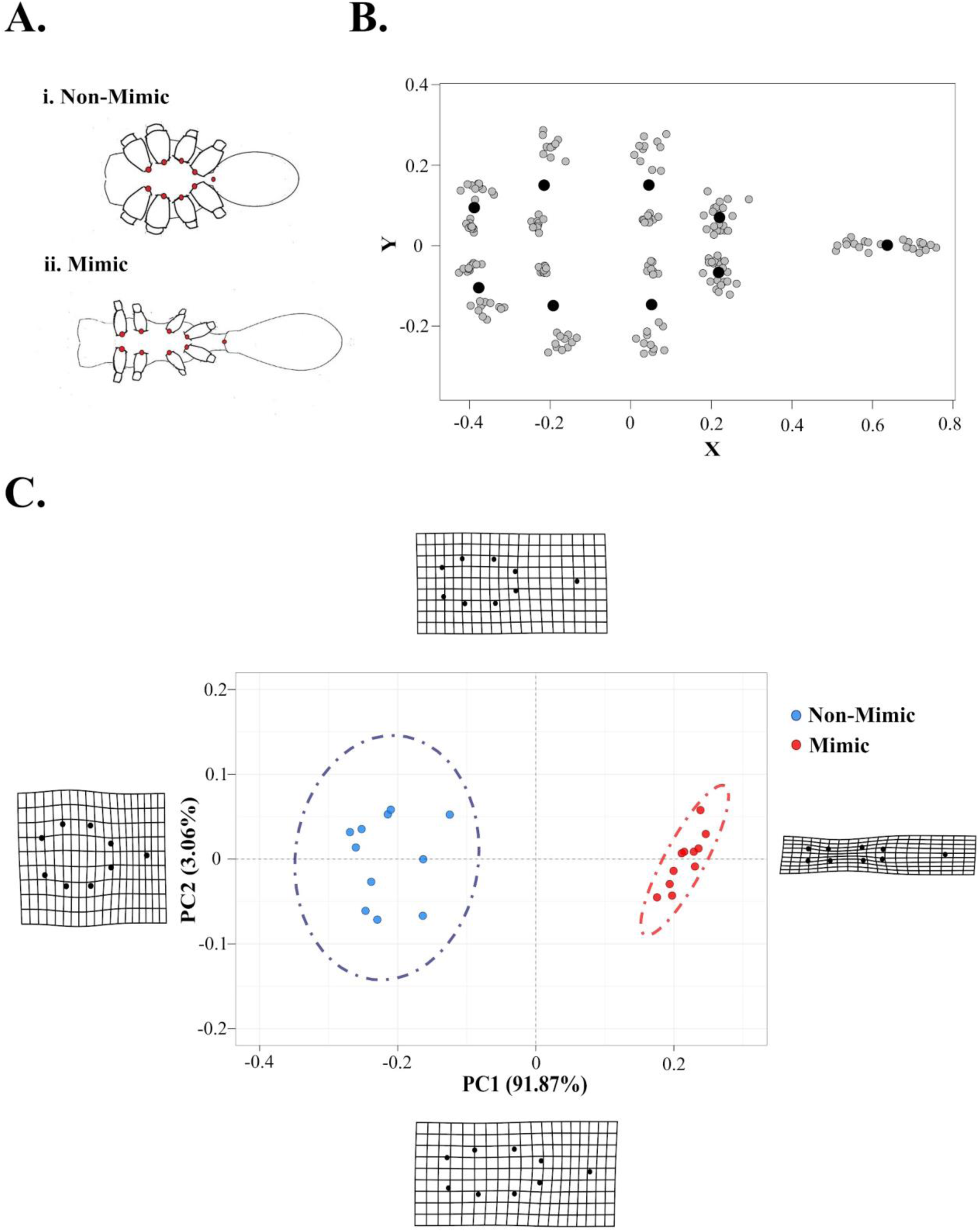
Shape variation in mimics and non-mimics. (A) Red dots represent the nine landmarks placed on the ventral side of i) non-mimics and ii) mimics. (B) Procrustes analysis of the landmarks is shown where grey points represent individual landmarks for each specimen, and the black points represent the shape consensus. (C) PCA based relative warp analysis shows non-mimics and mimics as two distinct groups along PC1 which explains 91.87% of the observed shape variation. Each data point represents the landmark configuration of an individual. The deformation grids shown on the sides of the PC plot represent the boundaries of the possible variation in shape.

We performed semilandmark based analysis to compare the body contour of the three groups. Model ants were included in this analysis as there is no requirement for homology when comparing semilandmarks. We imaged the dorsal side of the non-mimics (*Plexippus* sp.), mimics and model ants (N=10 for each), placed 120 equidistant points along the outline of their body (Fig. 5A), digitised these using *tpsDig* ver. 2.31 (Rohlf 2015, 2017) and superimposed them using *geomorph* (Adams & Otárola-Castillo 2013) (Fig. 5B). Similar to the landmark analysis, we performed relative warp analysis to compare the differences in the shape of the superimposed configurations.

**Fig.5.**
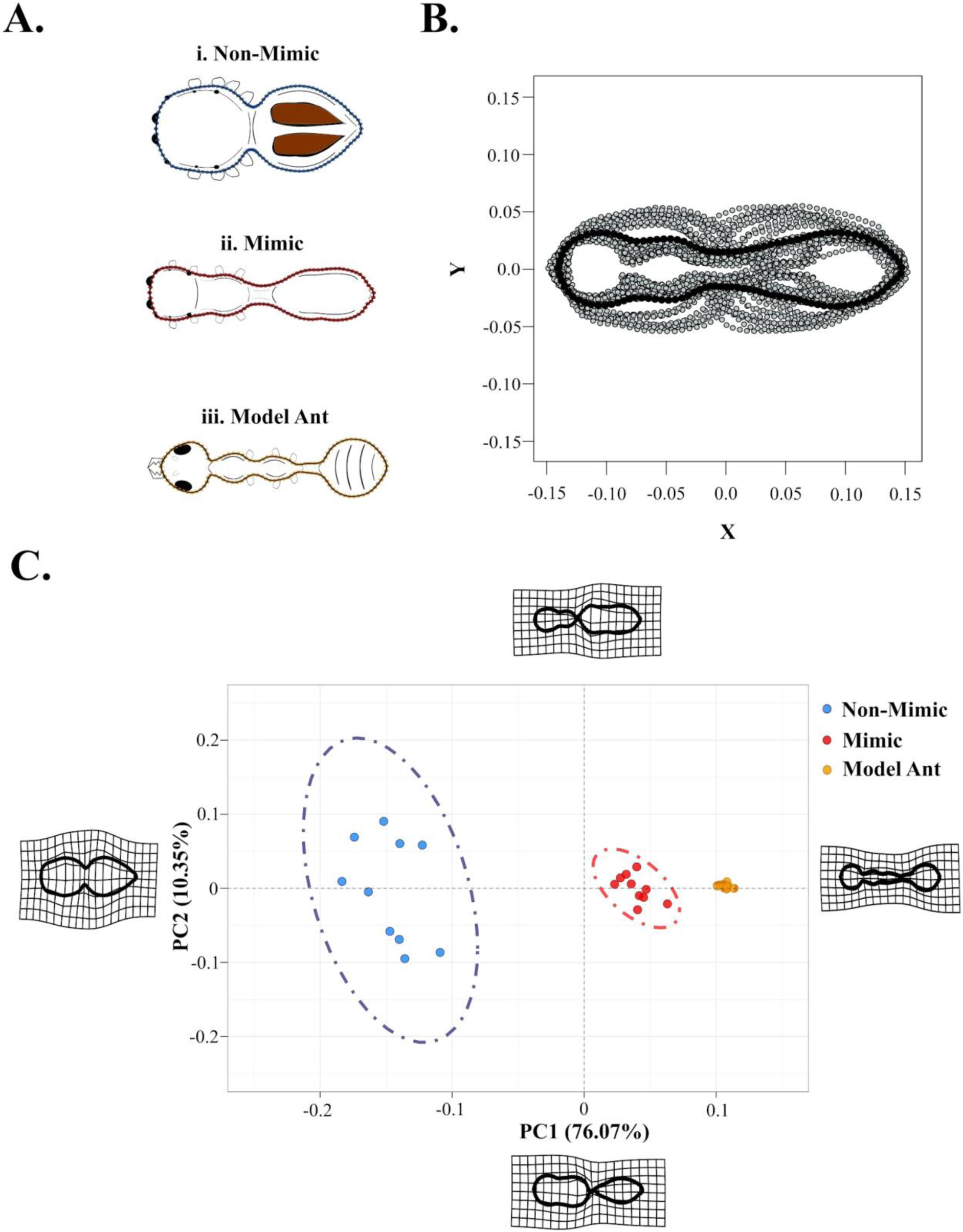
Differences in body contours of non-mimics, mimics and model ants. (A) 120 semilandmarks (coloured dots) along the body contours of (i) non-mimics, (ii) mimics and (iii) model ants. (B) Procrustes analysis of the semilandmarks is shown where grey points represent individual semilandmarks for each specimen, and the black points represent the shape consensus. (C) PCA based relative warp analysis of body contours shows mimics occupying an intermediate position between model ants and non-mimics along PC1 which explains 76.07% of observed shape variation. Here, each data point represents the body contour of an individual. The deformation grids shown on the sides of the PC plot represent the boundaries of the possible variation in shape.

## Results

### a) Speed of movement

Mimics moved at a speed that was comparable to that of model ants on a trail (Fig. 2). The speed of non-mimics (*P. petersi*) was approx. 385.5 times slower than the mimics. Speed of mimics (median = 0.0906 m/s) was comparable to that of model ants on trail (median = 0.0992 m/s) and significantly different from that of non-mimics (median = 0.000235 m/s) and model ants off trail (median = 0.236 m/s) (Fig. 2. Kruskal Wallis χ2 = 40.132, df = 3, p< 0.001; Dunn’s posthoc for pairwise comparisons: mimic vs. non-mimic p<0.005; mimic vs. ants on trail p=1, ants off-trail p<0.001). Ants on and off trail also showed significant differences in their speeds (p=0.005).

### b) Leg size

We present two sets of comparisons in this section - one between mimics and non-mimics (where we compare 5 leg appendages), and the other between non-mimics, mimics and model ants (where we compare only 3 common appendages). For each comparison, we present results for the right foreleg and hindleg. The results for left and right legs were similar. For mimics and non-mimics, the results of the remaining 3 pairs of legs are presented in the supplementary figure S1. For non-mimics, mimics and model ants, the results of the left leg appendages are presented in supplementary figure S2.

When we compared sizes of leg appendages of mimics to non-mimics, we found that they formed two distinct clusters in the morphospace along PC1 for both forelegs (PC1 88.68% explained variation) and hindlegs (PC1 74.06% explained variation) (Fig. 3B). All five leg appendages contributed equally to variation along PC1. Interestingly, there was more variation along PC2 for the hindlegs compared to the forelegs, as can be seen from the size of the ellipse and the percentage variation (18.09%). Tarsus contributed to most of this observed variation in the hindleg (Tarsus explained 94.78% of the variation in PC2).

When we compared non-mimics, mimics, and ants, we found that they form three distinct clusters along PC1 for both forelegs (80.27% explained variation) and hindlegs (93.12% explained variation) (Fig. 3C). All three appendages compared contributed similarly to variation along PC1. Along the PC2 axis, non-mimics and model ants are not distinct from each other (as their ellipses overlap), whereas the mimics are distinct from both these groups. This is especially apparent in the forelegs as can be observed by the size of the ellipse and the percent variation (17.9%). Tibia contributed to most of this observed variation in the foreleg (52.58% of the variation in PC2).

Mimics showed higher intra-specific variation in forelegs as evident by the larger size of the ellipse compared to the hindlegs (Fig 3C). In both forelegs and hindlegs, the variation observed in non-mimics was low despite including data from various genera. Ants also showed relatively high intraspecific variation even though only major workers were used in the study. No difference was observed between the leg size of ants belonging to two different colonies.

### c) Shape analysis

We compared shapes of non-mimics and mimics in this section. Landmark based relative warp analysis revealed that they formed distinct clusters along the PC1 axis (Fig.4C, 91.87% explained variance). The transformation warp-grids showed that the shape of non-mimics is oval compared to the streamlined, elongated shapes of mimics. The shapes encompassed by the mimics and non-mimics represent distinct body shapes that range from streamlined to oval.

When we compared the body contours using semilandmarks for non-mimics, mimics and model ants, they formed distinct clusters where PC1 accounted for 76.07% of the explained variance. Mimics were closer to the model ants than to non-mimics in the morphospace (Fig.5C). We found that ants showed relatively no variation compared to mimics and non-mimics and that the intraspecific variation was larger in non-mimics than mimics and model ants. Data from one colony of ants was used in this analysis.

## Discussion

Our results indicate that the mimics occupy an intermediate morphological space compared to the non-mimicking jumping spiders and model ants. This trend is true for both linear morphological measures of leg length and width, and for shape. This suggests that mimics face a unique set of challenges as they transform to match ants which are morphologically and phylogenetically distinct from spiders. On the other hand, we find that the speed of mimics is comparable to that of trail following ants. Thus, ant-mimicking spiders are better at mimicking some aspects of ant behaviour (in this case movement), than they are at mimicking morphology and overall body shape. This further raises questions about the limits to morphological change possible within these jumping spiders.

We find that the median speed of mimics is comparable to that of pheromone-trail following model ants, both of which were significantly higher compared to the speed of non-mimic *P. petersi* (Fig. 2). However, it is interesting to note that the speed of mimics was lower than that of model ants that do not follow pheromone trails. In nature, *O. smaragdina* worker ants rarely leave trails, and the trail following ants are much more aggressive than individual foragers (*pers obs*). It is therefore likely that mimicking the speed of a trail-following ant is advantageous. The movement of the mimics is in stark contrast to the movement shown by the non-mimetic jumping spider *P. petersi*, which is characterized by short bursts of walks and jumps interspersed with long stationary periods. Our results corroborate data from other systems that suggest that the movement of mimics is comparable to that of pheromone-trail following ants (Shamble et al. 2017), even for inaccurate mimics, and that such behaviour may be adaptive (Pekár & Jarab 2011).

Traditional morphometric analyses reveal morphological differences between ants, mimics and non-mimics, with the mimics occupying intermediate morphological space compared to the other two groups. Previous studies have either used a single measurement for the entire leg (Pekár et al. 2020), or have included only one part (femur) (Kelly et al 2021). Thus, our study adds depth to our understanding of the specific leg appendages that drive adaptations of mimics. All five leg appendages contribute equally to differentiating between mimics and non-mimics. However, along PC2, tarsus explained additional variation in the hindlegs. When we examined mimics, non-mimics, and model ants, we found that femur, tibia and tarsus contributed similarly to differentiating between the groups. However, along PC2, tibia contributed to additional variation in the foreleg. Tibia and tarsus are important hindleg appendages that contribute to walking and jumping in salticids (Hill 2018).

Phylogenetic and mechanical constraints may be imposing limits on the extent to which the leg morphology of the mimics can change, resulting in the observed intermediate morphology of mimics. Moreover, it is also possible that such incomplete morphological mimicry is sufficient to deceive predators, as predators often operate on insufficient information gleaned from potential prey (McLean & Herberstein 2021). This assertion is supported by our observation that mimics match the movement of ants quite well.

We find that spiders (both mimics and non-mimics) show lower intragroup variation for hindleg morphology when compared to ants. Further, for mimics, forelegs showed higher intraspecific variation than the hindlegs, much of which was explained by variation in tibia. Forelegs and hindlegs differ in their functions, with forelegs predominantly used for walking, and ‘antennal illusion’, and hindlegs for walking, and jumping (Hashimoto et al. 2020). Thus, it is possible that various selection pressures act on the different leg appendages. The mimics’ ability to jump and capture prey is also constricted by a slender body and constricted abdomen. Such differential selection has been reported in mimetic butterflies (Owens et al. 2020), where biomechanical requirements drove forewing morphology, and predation and sexual selection drove hindwing morphology. Interestingly, non-mimics showed very low variation in leg morphology even when individuals from multiple genera were included. The intraspecific variation observed for both colonies of weaver ants could emerge from body form differences within the same caste of ants, and size allometry (Tschinkel et al. 2003, Sommer & Wehner 2012).

Our analyses of landmarks and semilandmarks show that the mimics, non-mimics, and ants form three non-overlapping clusters in the morphospace. This pattern has been recovered in other studies that have also compared the shape of mimics and non-mimics (Hashimoto et al. 2020, Pekár et al. 2020). Further, in the morphospace defined by these shape parameters, mimics fall intermediate between model ants and non-mimics. Visually, although the mimics are very similar to model ants, quantitative assessments of their shape show that they occupy a distinct morphospace.

Our study focuses on comparing movement and form of mimics, non-mimics and model ants. Mimics move at a speed that is comparable to that of trail following model ants. On the other hand, mimics and model ants are distinct in the morphological space they occupy. This is true for both size and shape measurements. Mimicry is driven by the interplay of multiple traits; the relative contribution of these traits to mimetic resemblance is determined by both selection pressure by predators and evolutionary constraints. In a recent study, Pekár et al. (2020) compared movement, shape, size and colouration of mimics, non-mimics, and ants. Their results show that while ants and mimics overlap with respect to colouration and movement, they do not overlap when shape and size were compared. Taken with our results, these findings suggest that selection pressure for movement and coloration may be higher than for shape and size of the mimics. Such differential selection, possibly driven by predators operating with limited sensory information, allows for inaccurate mimics to persist in nature.

Selection pressures can also vary across the lifetime of an individual (Cushing 1997, Pekar et al. 2020). The interplay of these various selection pressures (including developmental pressures, sexual selection, among others) along with investigations into multiple traits is needed to understand the evolution of myrmecomorphy in spiders.

## Acknowledgements

Azim Premji University provided funding and logistical support to carry out this project. We would like to thank Pranav Balasubramanian who collected non-mimics used for this study. Part of the data collection and analysis happened during the pandemic. We would like to thank all our friends and family members for their support.

## References

Adams, D. C., & Otárola-Castillo, E. (2013). geomorph: an R package for the collection and analysis of geometric morphometric shape data. Methods in Ecology and Evolution, 4(4), 393–399.

Bates, H. W. (1862). XXXII. Contributions to an insect fauna of the Amazon Valley. Lepidoptera: Heliconidæ. Transactions of the Linnean Society of London, (3), 495–566.

Brown, D. (2008). Tracker Video Analysis and Modeling Tool. Version 5.1.4. http://physlets.org/tracker/

Ceccarelli, F. S. (2008). Behavioral mimicry in Myrmarachne species (Araneae, Salticidae) from North Queensland, Australia. The Journal of Arachnology, 36(2), 344–351.

Cushing, P. E. (1997). Myrmecomorphy and myrmecophily in spiders: a review. Florida Entomologist, 165–193.

Durkee, C. A., Weiss, M. R., & Uma, D. B. (2011). Ant mimicry lessens predation on a North American jumping spider by larger salticid spiders. Environmental Entomology, 40(5), 1223–1231.

Edmunds, M. (2006). Do Malaysian Myrmarachne associate with particular species of ant? Biological Journal of the Linnean Society, 88(4), 645–653.

Hashimoto, Y., Endo, T., Yamasaki, T., Hyodo, F., & Itioka, T. (2020). Constraints on the jumping and prey-capture abilities of ant-mimicking spiders (Salticidae, Salticinae, Myrmarachne). Scientific reports, 10(1), 1–11.

Hill, D. E. (2018). The jumping behavior of jumping spiders: a review (Araneae: Salticidae). Peckhamia, 167(1), 1–8.

Huang, J. N., Cheng, R. C., Li, D., & Tso, I.M. (2011). Salticid predation as one potential driving force of ant mimicry in jumping spiders. Proceedings of the Royal Society B: Biological Sciences, 278(1710), 1356–1364.

Kelly, M. B., McLean, D. J., Wild, Z. K., & Herberstein, M. E. (2021). Measuring mimicry: methods for quantifying visual similarity. Animal Behaviour, 178, 115–126.

Kropf, C. (2013). Hydraulic system of locomotion. In Spider ecophysiology (pp. 43–56). Springer, Berlin, Heidelberg.

Lawing, A. M., & Polly, P. D. (2010). Geometric morphometrics: recent applications to the study of evolution and development. Journal of Zoology, 280(1), 1–7.

Lê, S., Josse, J., Husson, F. & Mazet, J. (2008). “FactoMineR: A Package for Multivariate Analysis.” Journal of Statistical Software, 25(1), 1–18. doi: 10.18637/jss.v025.i01.

Mclver, J. D., & Stonedahl, G. (1993). Myrmecomorphy: morphological and behavioral mimicry of ants. Annual Review of Entomology, 38(1), 351–377.

McLean, D.J., & Skowron Volponi, M. A. (2018) trajr: An R package for characterisation of animal trajectories, Ethology 124, 440–448.

McLean, D. J., Cassis, G., Kikuchi, D. W., Giribet, G., & Herberstein, M. E. (2019). Insincere flattery? Understanding the evolution of imperfect deceptive mimicry. The Quarterly Review of Biology, 94(4), 395–415.

McLean, D. J., & Herberstein, M. E. (2021). Mimicry in motion and morphology: do information limitation, trade-offs or compensation relax selection for mimetic accuracy?. Proceedings of the Royal Society B, 288(1952), 20210815.

Nelson, X. J., Jackson, R. R., Edwards, G. B., & Barrion, A. T. (2005). Living with the enemy: jumping spiders that mimic weaver ants. The Journal of Arachnology, 33(3), 813–819.

Nelson, X. J., & Jackson, R. R. (2006). Vision-based innate aversion to ants and ant mimics. Behavioral Ecology, 17(4), 676–681.

Nelson, X. J., & Card, A. (2016). Locomotory mimicry in ant-like spiders. Behavioral Ecology, 27(3), 700–707.

Owens, H. L., Lewis, D. S., Condamine, F. L., Kawahara, A. Y., & Guralnick, R. P. (2020). Comparative phylogenetics of Papilio butterfly wing shape and size demonstrates independent hindwing and forewing evolution. Systematic biology, 69(5), 813–819.

Pekár, S., & KŘÁL, J. (2002). Mimicry complex in two central European zodariid spiders (Araneae: Zodariidae): how Zodarion deceives ants. Biological Journal of the Linnean Society, 75(4), 517–532.

Pekár, S., & Jarab, M. (2011). Life-history constraints in inaccurate Batesian myrmecomorphic spiders (Araneae: Corinnidae, Gnaphosidae). European Journal of Entomology, 108(2).

Pekár, S., Petráková, L., Bulbert, M. W., Whiting, M. J., & Herberstein, M. E. (2017). The golden mimicry complex uses a wide spectrum of defence to deter a community of predators. Elife, 6, e22089.

Pekár, S., Tsai, Y. Y., & Michalko, R. (2020). Transformational mimicry in a myrmecomorphic spider. The American Naturalist, 196(2), 216–226.

Pfennig, D. W., Harcombe, W. R., & Pfennig, K. S. (2001). Frequency-dependent Batesian mimicry. Nature, 410(6826), 323–323.

R Core Team (2021). R: A language and environment for statistical computing. R Foundation for Statistical Computing, Vienna, Austria. https://www.R-project.org/.

Ramesh, A., Vijayan, S., Sreedharan, S., Somanathan, H., & Uma, D. (2016). Similar yet different: differential response of a praying mantis to ant-mimicking spiders. Biological Journal of the Linnean Society, 119(1), 158–165.

Regier, J. C., Shultz, J. W., & Kambic, R. E. (2005). Pancrustacean phylogeny: hexapods are terrestrial crustaceans and maxillopods are not monophyletic. Proceedings of the Royal Society B: Biological Sciences, 272(1561), 395–401.

Rohlf, F. J., & Slice, D. (1990). Extensions of the Procrustes method for the optimal superimposition of landmarks. Systematic biology, 39(1), 40–59.

Rohlf, F. J. (2015). The tps series of software. Hystrix, 26(1).

Rohlf, F. J. (2017). tpsDig, version 2.31. Department of Ecology and Evolution, State University of New York at Stony Brook, Stony Brook, NY.

Rowe, C., & Halpin, C. (2013). Why are warning displays multimodal?. Behavioral Ecology and Sociobiology, 67(9), 1425–1439.

Ruxton, G. D., Allen, W. L., Sherratt, T. N., & Speed, M. P. (2004). Avoiding attack: the evolutionary ecology of crypsis, aposematism, and mimicry. Oxford University Press.

Schneider, C. A., Rasband, W. S., & Eliceiri, K. W. (2012). NIH Image to ImageJ: 25 years of image analysis. Nature Methods, 9(7), 671–675. doi:10.1038/nmeth.2089

Shamble, P. S., Hoy, R. R., Cohen, I., & Beatus, T. (2017). Walking like an ant: a quantitative and experimental approach to understanding locomotor mimicry in the jumping spider Myrmarachne formicaria. Proceedings of the Royal Society B: Biological Sciences, 284(1858), 20170308.

Sommer, S., & Wehner, R. (2012). Leg allometry in ants: extreme long-leggedness in thermophilic species. Arthropod structure & development, 41(1), 71–77.

Tschinkel, W. R., Mikheyev, A. S., & Storz, S. R. (2003). Allometry of workers of the fire ant, Solenopsis invicta. Journal of Insect Science, 3(1).

Uma, D., Durkee, C., Herzner, G., & Weiss, M. (2013). Double deception: ant-mimicking spiders elude both visually-and chemically-oriented predators. PLoS One, 8(11), e79660.

Vijayan, S., Balasubramanian, P., Casiker, C., & Uma, D. (2021). Non-mimetic jumping spider responses towards three species of ants and their mimics. Journal of Ethology, 39(1), 65–72.

