## Supplementary information for "An arachnid’s guide to be an ant: morphological and behavioural mimicry in ant-mimicking spiders"

**Supplementary Material**

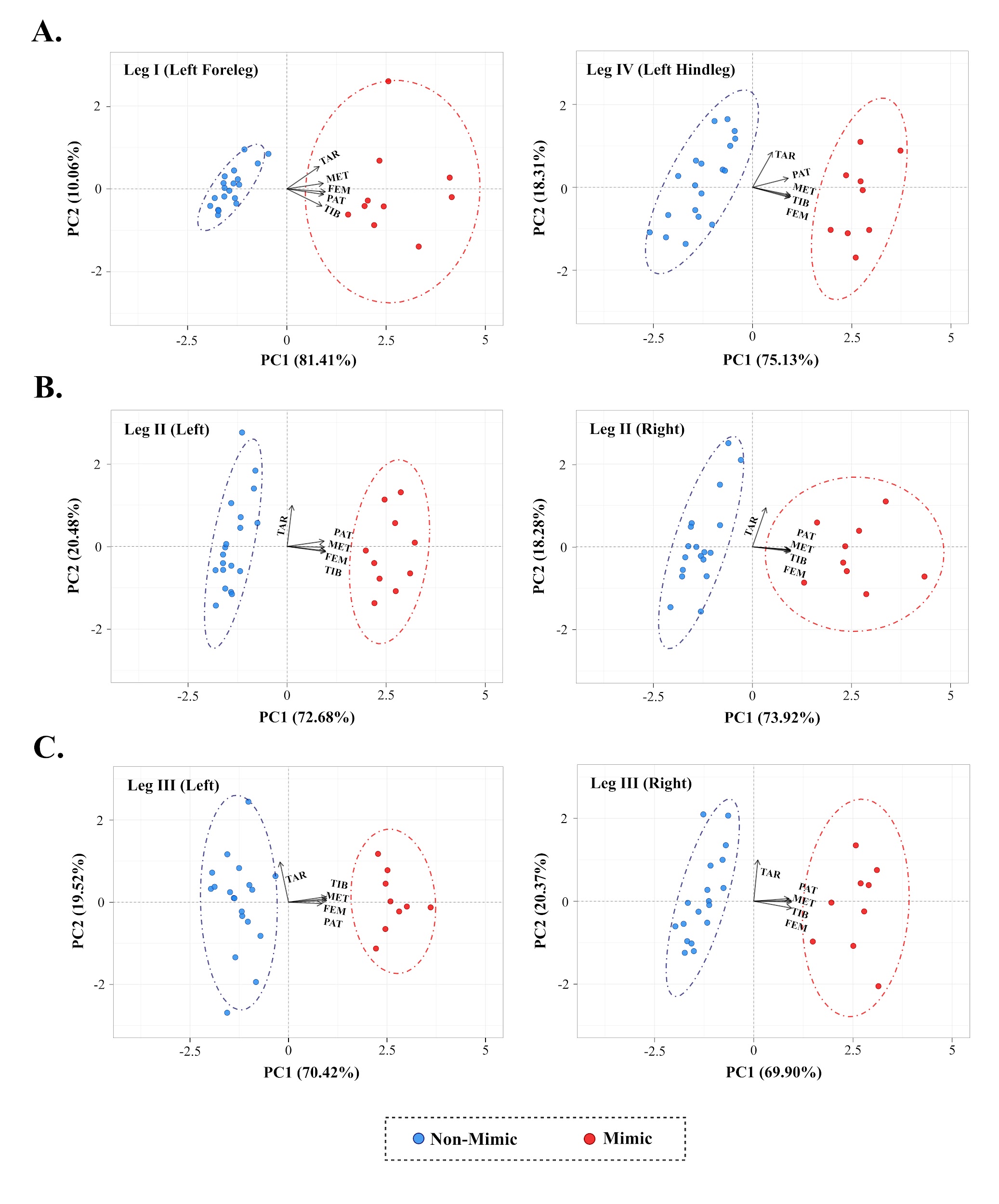

Fig.S1. PCA of leg size of non-mimics and mimics. (A), (B) and (C) depict PCA results quantifying size differences in the other six legs of non-mimics and mimics based on the length: width ratio of femur (FEM), patella (PAT), tibia (TIB), metatarsus (MET) and tarsus (TAR). For each leg, non-mimics and mimics show clear separation along the PC1 axis which accounts for the majority of the observed variation in leg size. Leg I = Foreleg and Leg IV = Hindleg. Each data point represents measurements taken from the leg of an individual.

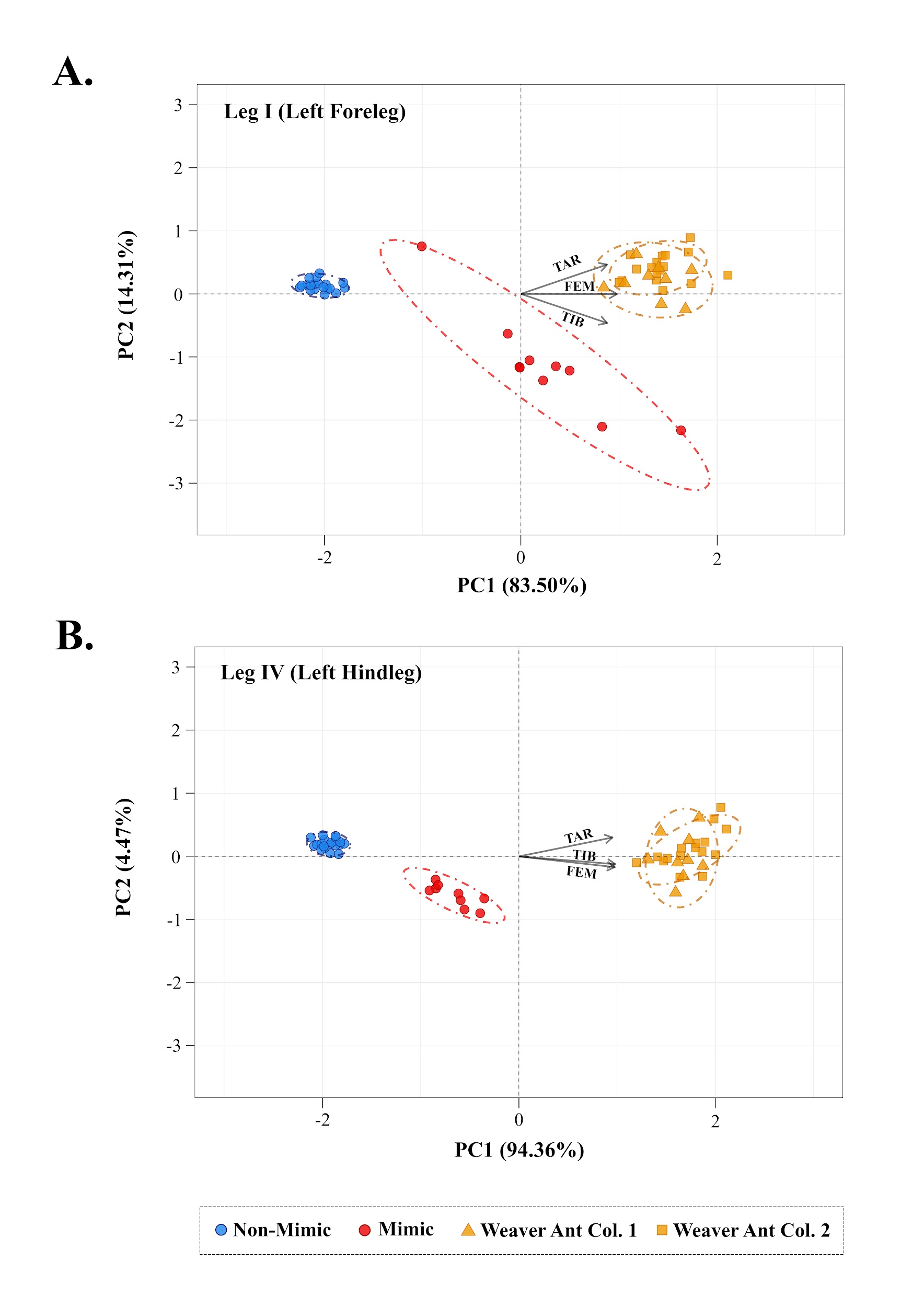

Fig S2. PCA of leg size of non-mimics, mimics and model ants. (A) and (B) show PCA results based on length: width ratio of leg appendages common to non-mimics, mimics and model ants viz. femur (FEM), tibia (TIB) and tarsus (TAR). Mimics occupy an intermediate position between non-mimics and model ants along the PC1 axis. PC1 explains 83.5% and 94.36% of observed variation in the size of left foreleg and hindleg respectively.

Table S1 - Contribution of different leg appendages to the first two PCs when comparing leg morphology of mimic and non-mimics. Leg I = Foreleg and Leg IV = Hindleg

| Appendage | Contribution to PC scores (in %) | | | | | | | | | | | | | | | | | | |
| --- | --- | --- | --- | --- | --- | --- | --- | --- | --- | --- | --- | --- | --- | --- | --- | --- | --- | --- | --- |
|  | Left | | | | | | | | | Right | | | | | | | | | |
|  | Leg I | | Leg II | | Leg III | | Leg IV | | | Leg 1 | | Leg II | | | Leg III | | | Leg IV | |
|  | PC1 | PC2 | PC1 | PC2 | PC1 | PC2 | PC1 | PC2 | PC1 | | PC2 | PC1 | PC2 | PC1 | | PC2 | PC1 | | PC2 |
| Femur | 22.99 | 1.00 | 25.59 | 1.08 | 26.51 | 0.17 | 23.13 | 6.59 | 21.34 | | 3.82 | 24.95 | 1.36 | 25.89 | | 2.73 | 25.08 | | 1.44 |
| Patella | 21.00 | 3.66 | 23.89 | 1.61 | 20.22 | 0.13 | 21.54 | 5.25 | 20.52 | | 3.79 | 22.58 | 0.50 | 22.52 | | 0.39 | 20.75 | | 0.01 |
| Tibia | 19.09 | 34.14 | 25.94 | 1.58 | 25.81 | 1.76 | 24.58 | 5.41 | 20.17 | | 4.31 | 24.05 | 0.64 | 25.91 | | 0.03 | 25.68 | | 2.88 |
| Metatarsus | 20.68 | 3.48 | 24.17 | 0.03 | 26.15 | 0.62 | 24.14 | 3.83 | 20.83 | | 5.29 | 25.30 | 0.73 | 25.38 | | 0.03 | 24.70 | | 0.89 |
| Tarsus | 16.24 | 57.71 | 0.42 | 95.70 | 1.32 | 97.32 | 6.62 | 78.92 | 17.15 | | 82.78 | 3.12 | 96.78 | 0.30 | | 96.83 | 3.78 | | 94.78 |

Table S2 - Contribution of different leg appendages to the first two PCs when comparing leg morphology of non-mimics, mimics and model ant. Leg I = Foreleg and Leg IV = Hindleg

| Appendage | Contribution to PC scores (in %) | | | | | | | |
| --- | --- | --- | --- | --- | --- | --- | --- | --- |
|  | Left | | | | Right | | | |
|  | Leg I | | Leg IV | | Leg I | | Leg IV | |
|  | PC1 | PC2 | PC1 | PC2 | PC1 | PC2 | PC1 | PC2 |
| Femur | 38.30 | 0.00 | 33.76 | 21.42 | 40.15 | 0.04 | 34.52 | 3.72 |
| Tibia | 30.90 | 49.68 | 34.12 | 11.27 | 29.39 | 52.58 | 33.32 | 33.72 |
| Tarsus | 30.80 | 50.32 | 32.13 | 67.31 | 30.46 | 47.38 | 32.16 | 62.56 |
